# Conformational changes and channel gating induced by CO_2_ binding to Connexin26

**DOI:** 10.1101/2020.08.11.243964

**Authors:** Deborah H. Brotherton, Christos G. Savva, Timothy J. Ragan, Victoria L. Linthwaite, Martin J. Cann, Nicholas Dale, Alexander D. Cameron

**Affiliations:** School of Life Sciences, University of Warwick, Gibbet Hill Road, Coventry, CV4 7AL, U.K; Leicester Institute of Structural and Chemical Biology, Department of Molecular and Cell Biology, University of Leicester, Lancaster Road, Leicester, LE1 7HB, UK; Department of Biosciences, Durham University, South Road, Durham DH1 3LE, UK; Biophysical Sciences Institute, Durham University, South Road, Durham DH1 3LE, UK

## Abstract

CO_2_ is the inevitable by-product of oxidative metabolism. Many physiological processes such as breathing^1^ and cerebral blood flow^2^ are sensitive to CO_2_. Historically, the physiological actions of CO_2_ have been regarded as being mediated exclusively *via* changes in pH. Here, we change this consensus by showing that the gap junction protein Connexin26 (Cx26) acts as a receptor for CO_2_ showing sensitivity to modest changes in PCO_2_ around the physiological norm^3-6^. Mass spectrometry analysis^7^ shows that CO_2_ carbamylates specific lysines on a regulatory loop of Cx26 at high, but not at low levels of PCO_2_. By means of high resolution cryo-EM, we have solved structures of Cx26 gap junctions at 1.9, 2.2 and 2.1 Å for PCO_2_ of 90, 55 and 20 mmHg respectively, all at pH 7.4. Classification of the particles at each level of PCO_2_, shows the transmembrane helices and N-terminal helix flexing at the dynamic cytoplasmic side of the protein. Gating of Cx26 gap junctions by CO_2_ involves movements of the N-terminus to plug the channel at high PCO_2_. We therefore provide mechanistic detail for a new paradigm by which CO_2_ can directly control breathing^8^ and other key physiological functions^9^.

## Introduction

There are 20 connexin genes in the human genome^10^. Connexins form hexameric plasma membrane channels, or hemichannels, that can dock together to form dodecameric gap junctions. Gap junctions provide a direct aqueous passageway between cells with functions independent of hemichannels^11-13^. Connexin mutations underlie many different pathologies affecting all organ systems of the body^14^. Mutations of Cx26 are a leading cause of congenital deafness^15^. Several rare, but dominant, mutations cause a severe syndromic disease: keratitis ichthyosis deafness syndrome (KIDS)^15^.

The conductance of gap junctions and hemichannels can be modulated by voltage^16,17^, pH^18-20^ and intracellular Ca^2+ 21^. Some connexins are modulated by NO^22,23^, and some by second messenger mediated phosphorylation^24^. We have recently discovered that a small group of closely-related *β*-connexins can be directly modulated by modest changes in physiologically relevant PCO_2_^3,4^. Physiological and mutational analysis suggests that this CO_2_ sensitivity is independent of pH^3,4^, intrinsic to the protein^4^, and mediated through a specific lysine by an unproven mechanism^3-5,25^. The CO_2_-sensitivity of Cx26 contributes nearly half of the centrally-generated chemosensory response to modest levels of hypercapnia^8,26^ and mediates CO_2_-dependent excitability changes in dopaminergic neurons of the substantia nigra and GABAergic neurons of the ventral tegmental area^9^.

The effect of CO_2_ on gap junctions and hemichannels is diametrically opposite: in hemichannels CO_2_ causes hemichannel opening^3-5,27^, whereas in gap junctions CO_2_ causes channel closure^6^. The closure of Cx26 gap junctions by a modest increase of PCO_2_ (*e.g.* from 35mmHg to 55 mmHg) can be mechanistically dissociated from the closing action of intracellular acidification. Mutation of Lys125 to an arginine in Cx26 gap junctions abolishes CO_2_ dependent closure, but has no effect on closure to intracellular acidification^6^.

In this paper, we provide a direct demonstration of the CO_2_-dependent carbamylation of purified Cx26 and suggest how this carbamylation reaction can mediate closure of the gap junction channel *via* consequent movement of the N-terminus.

### CO_2_ carbamylates purified Cx26

CO_2_ can bind to proteins through carbamylation reactions with neutral lysine *ε*-amino- or *N*-terminal *α*-amino-groups (Fig. 1). Such modifications affect the activities of RuBisCO^28^ and hemoglobin (Hb)^29,30^, respectively. Protein carbamylation can be directly detected by covalent trapping with the alkylation reagent triethyloxonium tetrafluoroborate (TEO)^7^. To investigate the presence of lysine carbamylation on Cx26, purified, DDM-solubilised Cx26 in buffer corresponding to a PCO_2_ of 90 mmHg and at a pH of 7.4, well below the unmodified pK_a_ of lysine, was treated with TEO to trap CO_2_^7^. This sample was digested with trypsin and analysed by ESI-MS-MS. Three residues were observed to have the additional mass associated with the alkylated carbamate group. This included Lys125 (Fig. 1e), which had been predicted to be the CO_2_-binding site, but also two other lysines on the same cytoplasmic loop (Lys108 and Lys122; Fig. 1d). Carbamates were identified on both peptides on internal lysine residues that exhibited a so-called missed cleavage. Carbamylation removes the cationic charge on the lysine that is essential for cleavage site recognition by trypsin resulting in such a missed cleavage. At a PCO_2_ of 20mmHg, also at pH 7.4, no carbamylation was observed (Fig. 1b-c; note that the figure shows similar peptides with missed cleavage but without the additional mass associated with the alkylated carbamate group). Cx26 therefore binds CO_2_ by lysine carbamylation and this carbamylation event is labile and sensitive to environmental PCO_2_. Further, the Cx26 lysine carbamylation profile is more complex than previous mutagenesis studies suggest. The carbamylation of these three lysine residues within the cytoplasmic loop, suggests that they share a common environment that favours this modification by CO_2_.

**Fig 1:**
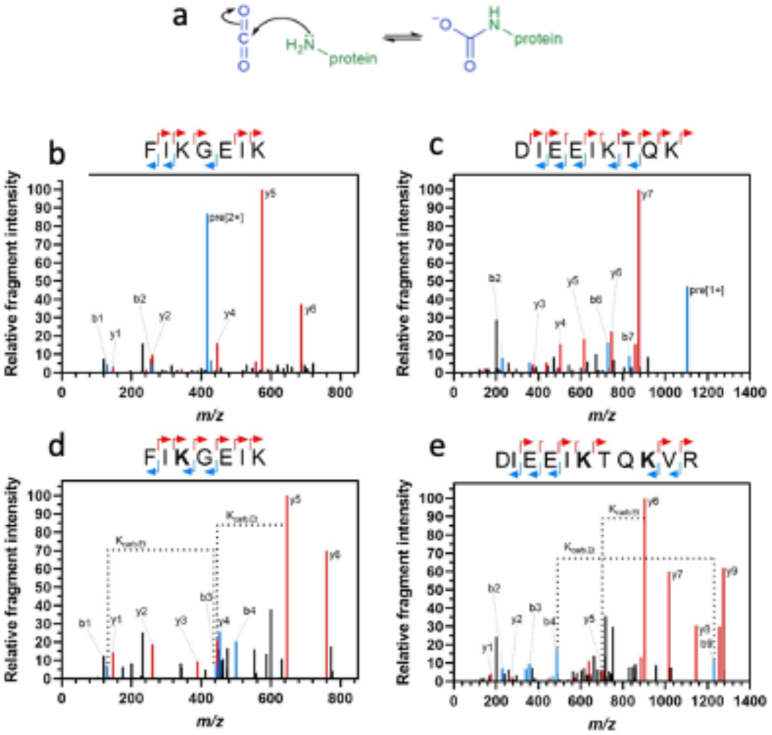
Identification of carbamate binding sites on hCx26 K108, K122 and K125. **a)** Reversible reaction of CO_2_ (blue) binding to a neutral amine on a protein (green). **b-e)** Plots of relative fragment intensity against mass/charge ratio (m/z) fragmentation data from ESI-MS-MS. Peptide sequences indicate the identification of y (red) and b (blue) ions by ESI-MS-MS. Under low total CO_2_/HCO_3_^-^ (20mmHg PCO_2_) conditions there is no carbamate identified on peptide K108 (**b**) or the peptide containing both K122 and K125 (**c**). Under high total CO_2_/HCO_3_^-^ (90mmHg PCO_2_) condition there are carbamates identified on K108 (**d**) and both K122 and K125 (**e**).

### Cryo-EM structure of Cx26 at 55mmHg PCO_2_

The structure of Cx26 was determined using cryo-EM from protein vitrified in a CO_2_/HCO_3_^-^ buffer corresponding to a PCO_2_ level of 55mmHg and a pH of 7.4. Under these conditions the gap junction would be expected to be largely closed^6^. 3D reconstructions were obtained at a nominal resolution of 2.5Å as defined by gold standard Fourier Shell Correlations^31,32^ (FSC) without applying symmetry and 2.2Å when D6 symmetry, consistent with the dodecameric structure of the gap junction, was used (Figure 2a; Extended Data Fig.1 and Extended Data Tables 1 and 2). Whereas similar reconstructions derived from similar protein vitrified in HEPES did not go beyond 4Å, the resolution of the structure under these conditions was limited only by the pixel size of the detector, suggesting that the CO_2_/HCO_3_^-^ buffer was stabilising the protein.

**Fig. 2:**
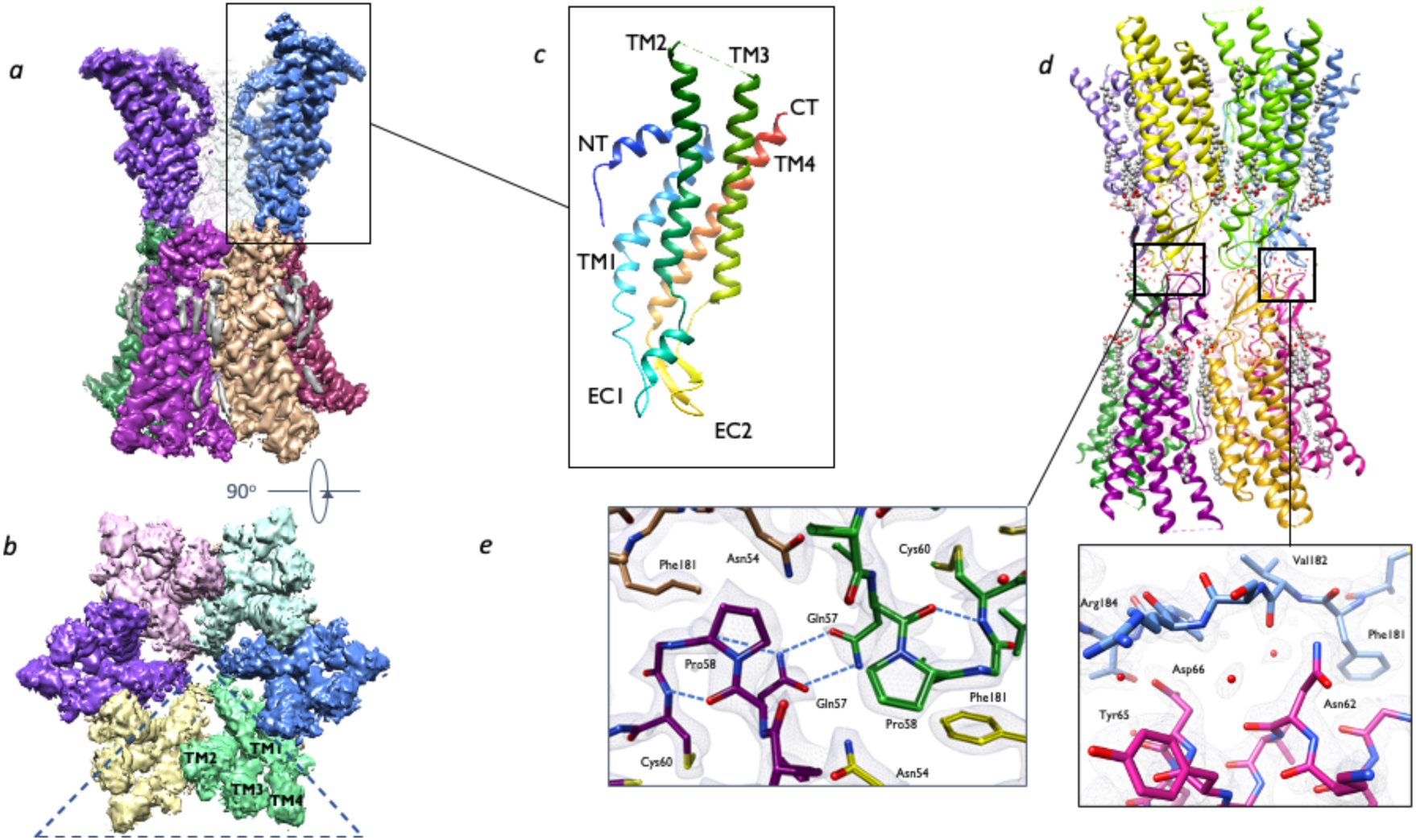
Structure of Cx26 solved by cryo-EM in CO_2_/HCO_3_^-^ buffer at 55mmHg PCO_2_. ***a-b)*** Cryo-EM Density for Cx26. The density has been coloured according to the associated subunit of the dodecameric gap junction with lipid/detergent chains in grey. **a)**: view perpendicular to the membrane, with two subunits and associated lipids removed to view the channel core. **b)**: view from the cytoplasmic side looking into the pore. The N-terminal helices fold into the pore to form a narrow restriction. **c)** The Cx26 subunit coloured from blue at the N-terminus to red at the C-terminus. **d)** Ribbon diagram corresponding to the density in (a). Water molecules and lipid/detergent are shown as spheres. The inset **(e)** shows the sharpened density in the region of the gap junction EC1 and EC2. Carbon atoms are coloured by chain.

The connexin subunit is composed of 4 transmembrane helices (TM1-4) with an N-terminal helix, an extracellular domain and a cytoplasmic loop^33^ (Fig. 2). Six connexin subunits form the hemichannel, which dock together to form the gap junction. We found excellent density in the gap junction region of the protein (Fig. 2; Extended Data Fig. 2). The high resolution of the maps has also allowed us to locate water molecules, many of which are instrumental in forming subunit interactions (Fig. 2). Lipids or detergents can be observed to bind in the dimples between the subunit interfaces (Fig. 2). On the other hand, the cytoplasmic side of the protein is less well defined with no consistent density for the cytoplasmic loop between TM2 and TM3 (Extended Data Fig. 2).

The resolution is much higher than has been seen for previous structures of Cx26^20,33,34^, enabling the protein to be modelled more accurately. Relative to these structures, the most interesting differences involve the N-terminus. The N-terminal helices fold into the pore such that the cryo-EM maps show a narrow constriction where the six helices meet (Fig. 2). This helix is known to be flexible and while there may be some ambiguity of the modelling of this region in both the crystal structure^35^ and in our structure (see methods), it is clear that the position of the helix and hence the shape of the pore entrance differs between structures (Fig. 3, Extended Data Fig. 3). In Cx46/50, the only other connexin structure that has been solved to date, the N-terminal helix has clear side-chain density^35,36^. In molecular dynamics simulations the N-terminal helix of Cx46-50 is more stable than its equivalent in the crystal structure of Cx26^35^. The conformation of the helix in our cryo-EM structure is more similar to that of Cx46/50 except that in the latter the helix is much more tucked back towards TM1 giving a more open pore (Fig. 3, Extended Data Fig. 3).

**Fig. 3:**
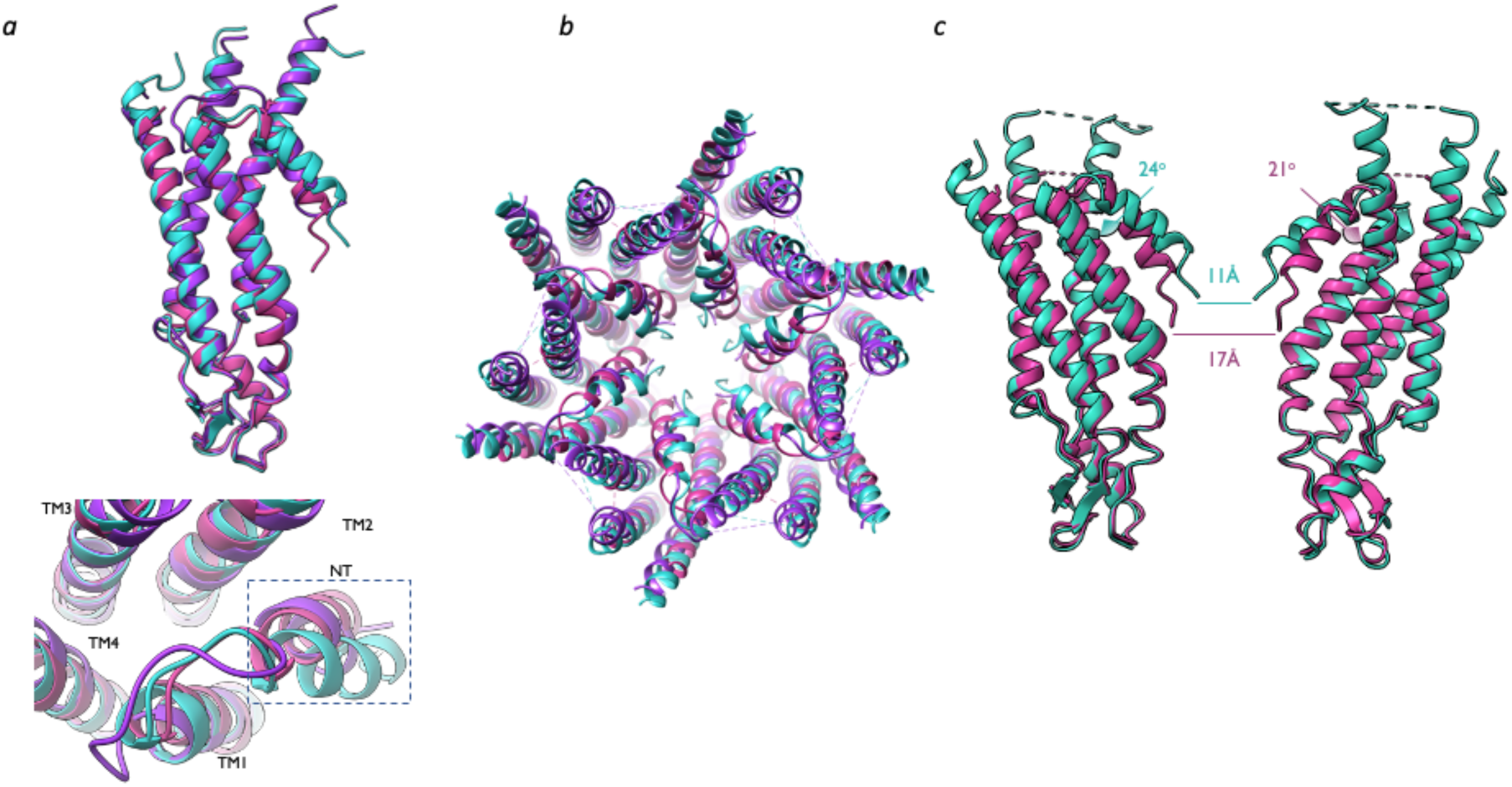
Comparison of cryo-EM Cx26 to other connexins. **a)** Superposition of Cx26 cryo-EM structure (cyan) on the crystal structure of Cx26 (2ZW3^33^; purple) and the cryo-EM structure of Cx46 (6MHQ^35^; violet). Superposition of one subunit of each protein, coloured as above (root mean square deviation (rmsd) cryo-EM Cx26 to 2ZW3: 1.0Å across 159 C_α_ pairs within 2Å after superposition; 2.1Å across all 189 C_α_ pairs, cryo-EM Cx26 to 6MHQ: 0.8Å across 152 C_α_ pairs within 2Å after superposition; 1.7Å across all 177 C_α_ pairs. Views from the side *(top)* and top *(bottom).* **b)** Superposition over the six subunits of the hemichannel (rmsd cryo-EM Cx26 to 2ZW3: 1.2Å across 834 C_α_ pairs within 2Å after superposition; 2.1Å across all 1002 C_α_ pairs, cryo-EM Cx26 to 6MHQ: 1.1Å across 869 C_α_ pairs within 2Å after superposition; 2.0Å across all 1062 C_α_ pairs). The protein is viewed from the cytoplasmic side of the membrane. **c)** Comparison of the angles of the N-terminal helices between the cryo-EM Cx26 and Cx46 structures. Two subunits are shown. The measurements shown are between corresponding C_α_ atoms. Subtle changes in modelling can reduce this.

### Conformational variability in Cx26

The cytoplasmic part of Cx26 is clearly flexible as judged, not only from the density and structural comparison, but also from elastic network modelling, where considerable flexing of the subunits and transmembrane helices can be observed^4^. As cryo-EM data sets can harbour a range of conformations, we analyzed particle subsets in two ways. First, we used variability analysis as implemented in cryoSPARC^37^ (Extended Data Movie 1). Clear movements can be observed in the N-terminus with the N-termini seeming to appear and disappear. Together with these movements there is a twisting of the cytoplasmic portions of the subunits. The greatest flexing of the TM helices is observed for TM2, which is located between the N-termini of two neighbouring subunits and appears to sway into and out of the channel (Extended Data Movie 1).

To obtain a more quantitative estimate of the range of conformations sampled by the protein we used a process of particle expansion, subtraction and masked classification in Relion^32,38^. When masks were created around a single subunit or a hemichannel it was difficult to discriminate between the classes. However, classification was more effective when a mask covering two subunits was constructed (Fig. 4). Qualitatively, the result was similar to that obtained from the cryoSPARC analysis. The first notable feature was the quality of the density associated with the N-terminus and its linker to TM1 (Fig. 4). Models were refined against each of the reconstructions (Extended Data Table 3). Together with the N-terminus, TM2 showed the most variability, flexing around Pro87 followed by TM3 flexing around Arg143.

**Fig. 4:**
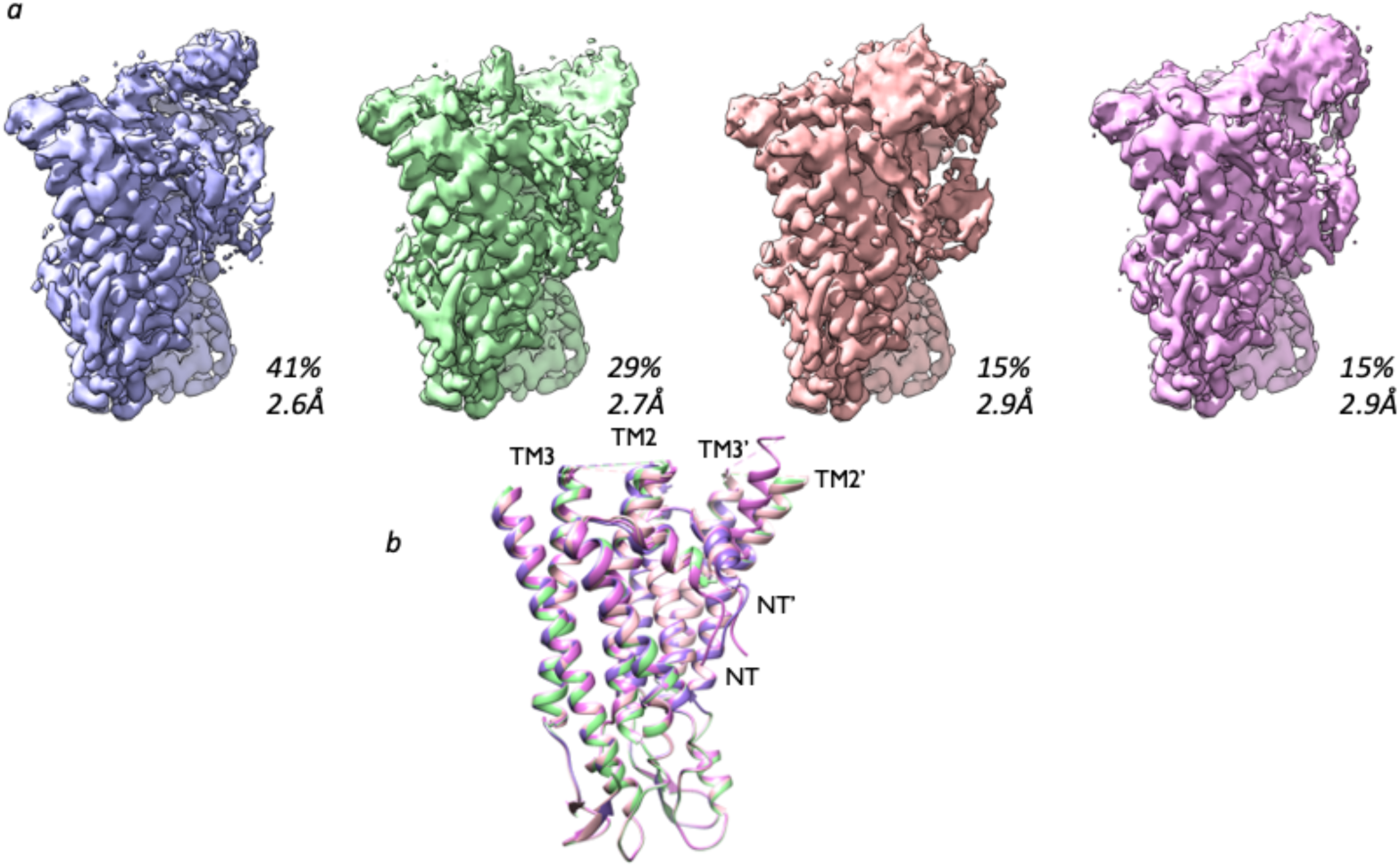
Variability in the structure of Cx26. **sa)** A two-subunit mask was used for particle subtraction and masked classification in Relion^32^. The data were split into four classes as shown. Particle reconstruction (see methods) resulted in maps with the given resolution. Generally, of the two N-termini, the density for one was better than the other and, in fact the best density was observed for one of the particle sets associated with only 15% of the particles. **b)** Adjustment of the model together with real-space refinement resulted in four models. These are coloured according to the associated map. Where the N-terminus could not be observed in the map, this was deleted. A morph between the pdb files of the two 15% classes can be seen in Extended Data Movie 2. In the class with the best-defined N-terminal helix, the two TM2s from the respective subunits are closer together than observed in the other classes.

### Comparison of structures associated with high and low CO_2_ levels at constant pH

To investigate the effect of CO_2_ on conformation further, we collected data from protein vitrified under levels of CO_2_ where the pore should be open (20mmHg) and a complementary data set at high CO_2_ (90mmHg), where it should be closed (hereafter respectively referred to as hypocapnic and hypercapnic CO_2_ structures). Both these data sets were collected from the same batch of protein and the buffers were chosen to ensure that the pH and ionic strength did not differ (see methods and Extended Data Fig. 4).

We obtained similarly high-resolution maps to the previous structure (1.9Å and 2.1Å with D6 symmetry for the hyper- and hypocapnic structures respectively and 2.1Å and 2.7Å, without symmetry averaging; Extended Data Figs 5-8, Extended Data Tables 1 and 2). The most striking difference apparent in the initial maps was the quality of the density for the N-terminal helix. While the density in this region of the maps associated with the hypercapnic structure was similar to the previous maps, that associated with the hypocapnic structure was much weaker giving the appearance of a more open pore (Fig. 5). This is consistent with our prior observations that hypercapnic conditions close Cx26 gap junctions by a mechanism independent of pH^6^. We used further classification procedures, with matched particle numbers to minimise artefacts caused by differences in processing, to investigate the effects of CO_2_. All classifications were consistent with a less defined helix for the N-terminus in the hypocapnic structure. Most interesting was the classification with a mask comprising 2 subunits as described above. Even though the classifications were not driven primarily by the N-terminal helix position, there was a difference in its associated density in each of the classes. While classification of the hypercapnic data gave the top class with 78% of the particles with well-defined N-terminal helices, the N-terminus in the top class of the hypocapnic data (58% of particles) was very ill defined (Extended data Fig. 9). It would appear, therefore that the higher CO_2_ levels bias and stabilize the dynamic equilibrium of the position of the N-terminal helix consistent with closing the channel.

**Fig. 5:**
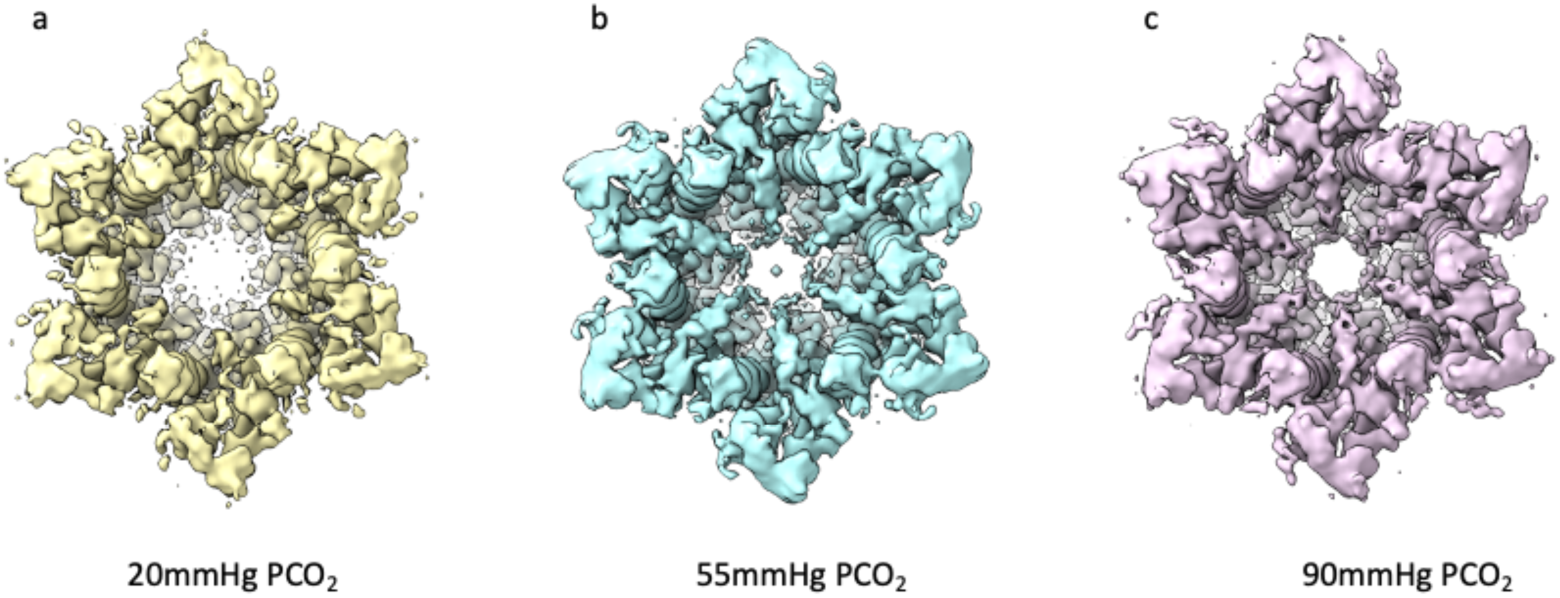
Variation in N-terminal density depending on levels of CO_2_. View from the cytoplasmic side of the channel. At low CO_2_, the density for the N-terminus is less defined given an apparently more open pore **a)** 20mmHg **b)** 55mmHg **c)** 90mmHg. Refinement and reconstructions were carried out using data sets of 125,000 particles with D6 symmetry. Maps have been low-pass filtered to 3Å to minimise noise and thresholds have been matched based on the density in the gap junction region.

### Mutations associated with KID syndrome

There are nine mutations that give rise to KIDS^15^ (Extended Data Movie 2). Four of these mutations occur in the highly flexible regions of the protein near the flexion point of TM2 and in the linker between the N-terminal helix and TM1. These mutations involve substitution of bulkier side chains (Ala88Val and Gly12Arg, Asn14Lys/Tyr and Ser17Phe), which would likely impede the flexions required for channel gating (Extended Data Movie 2). Ala88 is situated next to Pro87 at the flexion point of TM2, and interacts with Val13 on the N-terminal helix adjacent to Asn14. By contrast, Ala40 is on TM1, and its position in the cryo-EM structures differs from that in the crystal structures due to a change in the conformation of residues Val37 to Glu42 (Extended Data Figure 3). This region also appears to be flexible and an interplay between Lys41, Glu42 and the N-terminus alters voltage gating^39^. Glu42 and Glu47 also form a binding site for Ca^2+ 34^ which may be important for hemichannel closure. Whereas in the crystal structures, the replacement of Ala40 by the bulkier valine might be tolerated, there is less space at the position it adopts in the cryo-EM structure and this mutation would likely distort the positioning of the transmembrane helices (Extended Data Fig. 3) and give rise to altered channel gating^6,40,41^.

## Discussion

While we have previously demonstrated the CO_2_-sensitivity of Cx26 hemichannels and gap junctions, the mechanism by which CO_2_ alters channel gating had not been established^3,4,6,25^. Our results definitively demonstrate that the actions of CO_2_ involve carbamylation of specific lysine residues in the cytoplasmic loop of the protein. Lys125 is necessary for the CO_2_ sensitivity of the channel, and mutation of this residue to Arg destroys its CO_2_-sensitivity. However, we find that two other nearby lysine residues are also carbamylated: Lys108 and Lys122. Carbamylation of lysine requires a restricted hydration space to alter the pK_a_ of the *ε* amino group and ensure that is not protonated at physiological pH. A simple hypothesis is that Lys108, Lys122 and Lys125 are within a common chemical environment that permits their carbamylation.

The carbamylated lysine residues are located in a mobile loop between TM2 and TM3, the cytoplasmic ends of which are also flexible. Classification of particles indicates that the positions of TM2 and TM3 can influence the conformation of the N-terminal helix. Under hypercapnic conditions the N-terminus is much more defined than in a similar structure collected from protein under low CO_2_ conditions, where we have ensured the buffer is at the same pH. The gating action of CO_2_ on Cx26 is therefore mediated *via* movements of the N-terminal helix and its ability to plug the channel. This same mechanism (movement of the N-terminal helix) has been proposed to gate the channel in response to signals such as voltage or pH^20^. Presumably, the structural drivers and critical residues for initiating movement of the N-terminal helix are different for these three modifiers of Cx26 gating. Interestingly, whereas profound acidification is required to cause the N-terminal helix to plug the channel, suggesting a function only in serious pathology, our findings that modest changes in PCO_2_ can utilize this same gating mechanism showing that it can play a role in normal physiology too.

Cx26 imparts CO_2_-sensitivity to the control of breathing^8,26^ and causes CO_2_ dependent excitability on neurons of the substantia nigra and ventral tegmental areas of the brain^9^. The KIDS mutations of Cx26, Ala40Val, Ala88Val and Asn14Lys/Tyr, abolish CO_2_ sensitivity^41-43^. This may be important for human health, as infants with the Ala88Val mutation are subject to bouts of central apnoea^43,44^. Our structures of Cx26, at different levels of CO_2_, demonstrate how carbamylation can modulate vital physiological functions by acting as an important post-translational modifier of ion channel gating.

## Methods

### Protein expression and purification

Human connexin 26 protein (hCx26) was expressed with a thrombin-cleavable His(6) affinity tag on the C terminus using baculovirus in *Spodoptera frugiperda* (Sf9) cells. This construct was a gift from Prof Tomitake Tsukihara. Cells were harvested 72 hours post infection by centrifugation at 2500 x g in a Beckmann JLA 8.1000 rotor, cell pellets were snap frozen in liquid nitrogen, and stored at −80°C until purification. Cells were thawed in hypotonic lysis buffer, (10 mM sodium phosphate, 10 mM NaCl, 5 mM MgCl_2_, 1 mM DTT, pH 8.0) DNAse I, cOmplete™ EDTA-free Protease Inhibitor Cocktail (Roche) and AEBSF were added according to manufacturer’s instructions before use. After 30 minutes stirring at 4 °C, the cells were broken using 40 strokes with a dounce homogeniser, and the membranes separated by ultracentrifugation for 1 hour at 4 °C, 158000 x g. Membranes were resuspended in membrane resuspension buffer (25 mM sodium phosphate, 150 mM NaCl, 5 % glycerol, 1 mM DTT, pH 8.0). DNAse I, Complete protease inhibitors and AEBSF were added according to manufacturer’s instructions before use. This membrane suspension was diluted (to 400 ml) with solubilisation buffer (10 mM sodium phosphate, 300 mM NaCl, 5 % glycerol, 1 mM DTT, 1% DDM (Glycon Biochemicals GMBH), pH 8.0) and incubated at 4 °C for 3 hours, before a further 1 hour ultracentrifugation at 4 °C, 158000 x g to remove insoluble material. The soluble material was batch bound to pre-equilibrated HisPur Ni-NTA resin (Thermo Scientific) overnight, and then poured into an econocolumn for subsequent manual washing and elution steps. Resin was washed with 5x CV wash buffer (10 mM sodium phosphate, 500 mM NaCl, 10 mM histidine, 5 % glycerol, 1 mM DTT, 0.1 % DDM, pH 8.0) before eluting hCx26 with elution buffer (10 mM sodium phosphate, 500 mM NaCl, 200 mM histidine, 5 % glycerol, 1 mM DTT, 0.1 % DDM, pH 8.0). Fractions containing hCx26 were dialysed overnight 4 °C against (10 mM sodium phosphate, 500 mM NaCl, 5 % glycerol, 1 mM DTT, 0.03 % DDM, pH 8.0). For the protein used for electron microscopy thrombin (Sigma) 1:1 w/w was added during dialysis to remove the affinity label). hCx26 was then 0.2 μm filtered, concentrated in a vivaspin 100,000 MWCO and loaded onto a Superose 6 5/150 size exclusion chromatography column (GE Healthcare Lifescience) to remove thrombin and exchange the buffer to either 20 mm, 55 mm or 90 mmHg *α*CSF buffer^3^ (*20 mmHg αCSF buffer*: 140 mM NaCl, 5 % glycerol, 1 mM DTT, 0.03 % DDM, 10 mM NaHCO_3_, 1.25 mM NaH_2_PO_4_, 3 mM KCl, 1 mM MgSO_4_, 2 mM MgCl_2_; *55 mmHg αCSF buffer*: 100 mM NaCl, 5 % glycerol, 1 mM DTT, 0.03 % DDM, 50 mM NaHCO_3_, 1.25 mM NaH_2_PO_4_, 3 mM KCl, 1 mM MgSO_4_, 2 mM MgCl_2_; *90 mmHg αCSF buffer*: 70 mM NaCl, 5 % glycerol, 1 mM DTT, 0.03 % DDM, 80 mM NaHCO_3_, 1.25 mM NaH_2_PO_4_, 3 mM KCl, 1 mM MgSO_4_, 4 mM MgCl_2_).

### Cryo-EM sample preparation and data collection

The hCx26 peak was concentrated to 3.5 mg/ml before being gassed with the correct amount of CO_2_ to give a final pH of ∼7.4 as described below for each buffer. 2.5%, 10% and 15% CO_2_ in N_2_ (BOC) were used for the 20, 55 and 90mmHg buffers respectively. The pH was controlled in two steps: First, 10ml of buffer was equilibrated with the correct percentage of CO_2_ for the buffer and the pH checked using a pH meter. Phenol red solution (1:200) was added to these samples and the colour noted for both ungassed and equilibrated buffer. The volume of the protein to be used for vitrification was noted, and an equal volume of phenol red/buffer was added to two separate Eppendorf tubes, one of which was gassed until the colour matched an equal volume of the pre-gassed 10μl sample. The time of gassing was noted, and the protein was treated identically (Extended Data Fig. 4). For both the 20mmHg and 90mmHg conditions the sample was left overnight dialysing against 60ml buffer that was gassed with the correct concentration of CO_2_ to give a pH of 7.4. Quantifoil 300 mesh gold grids (either 0.6/1 carbon film, 0.6/1 or 1.2/1.3 UltrAUfoil, Quantifoil Micro Tools GMBH) were glow discharged for 30 seconds prior to use. Vitrification of the protein in liquid ethane/propane at −180 °C was carried out with a Leica GP2 automated plunge freezer with 3 μl protein per grid at 4 °C, 95 % humidity, 6 seconds blotting in a CO_2_ / N_2_ atmosphere appropriate for the buffer used. Grids were screened using a Jeol 2100plus microscope, and data were collected on an FEI Titan Krios G3 equipped with a K3 detector and BioQuantum energy filter using a 20 eV slit width. A dose rate of ∼10 e/pix/sec on the detector was chosen with a final dose of between 40-45 e/Å2 on the specimen. Data collections were prepared and run automatically using EPU2.X and aberration-free image shift (AFIS).

### Cryo-EM data processing

Data were processed using Relion3.1-beta^38^, using essentially the same protocol for the three data sets, accounting for the different pixel size. Micrographs were motion corrected using the version of MotionCor2^45^ implemented in Relion, and CTFs were estimated using ctffind4^46^. Particles were picked using the Laplacian of Gaussian (LoG) picker, and poor, damaged, or junk particles were removed by serial rounds of 2D classifications with particles downsampled to 4Å/pixel. 3D classification was carried out in C1 with an initial model generated from a previous Cx26 cryo-EM structure (unpublished data) with a low-pass filter of 30Å. Multiple rounds of 3D classification in C1 resulted in 4 very similar classes. Exhaustive rounds of refinement, CTF refinement and polishing in Relion with unbinned particles were used to improve the resolution of the Coulomb shells until no further improvement was gained. Trials were made treating the classes individually or pooling them in C1, C2, C3, C6 and D6 symmetry. While variations were noted in the associated maps, it was not possible to identify different structural features. The resolution was estimated based on the gold standard Fourier Shell Coefficient (FSC) criterion^31,32^ with a soft solvent mask. Local Resolution estimation was carried out in ResMap^47^.

### Variability Analysis in cryoSPARC

The 344085 refined particles associated with the 55mmHg data set were downsampled to 1.586Å/pixel and imported into cryoSPARC^37^. An ab-initio reconstruction was made in C1 and the particles refined in cryoSPARC using D6 symmetry to give maps with an estimated resolution of 3.24Å. Particle expansion was carried out with D6 symmetry. These were then subjected to variability analysis in cryoSPARC with a filter resolution of 4.5Å and a mask over the six subunits of one of the two docked hemichannels that form the gap junction. The results were displayed as a simple movie of 20 frames shown in Chimera^48,49^. Analysis was also carried out with masks covering single or neighbouring subunits, but these did not appear to give any advantage over the hemichannel mask.

### Particle subtraction and masked classification in Relion

For consistency among the three data sets, which contain different numbers of particles, 125,000 particles were randomly selected from each of the 55mmHg and 90mmHg CO_2_ D6 refined particle sets to match the particle numbers in the 20mmHg CO_2_ data set. These were then symmetry expanded in D6. A mask was created in Chimera^48,49^ based on two neighbouring subunits of the refined model and a soft edge added in relion_mask_create. This was then used in particle subtraction and 3D classification without image alignment. Reconstructions were created with half subsets of each class and the resolution was estimate with relion_postprocess.

### Model Building

Initial model building was carried out in Coot^50^ with the D6 reconstructions of the 55mmHg CO_2_ data with an initial model based on the crystal structure of Cx26^33^. Residues at the C-terminus and the cytoplasmic loop between 102 and 128 could not be observed and were not included in the model. While the density clearly showed the position of the N-terminal helix the side chains were not well defined and modelling of the first residues is ambiguous. For this reason we have omitted the first three residues in the final PDB file. Real space refinement in Phenix^51^ was carried out with NCS constraints. Water molecules were also added during refinement with further curation of the water molecules added in Coot. Lipids or detergents were clearly visible in the density, though the exact nature of the head groups is ambiguous. The head-groups have been omitted in the final structure. Refinement of the 20mmHg and 90mmHg structures was carried out starting from a partially refined structure of the 55mmHg CO_2_ structure using a similar protocol. The structures of the four classes of the 55mmHg CO_2_ data set were also refined in a similar manner.

### Structural Analysis

All structural images shown in this paper were generated in Chimera^48,49^ or Chimera X^52^. Superpositions were carried out in Chimera such that only matching C_α_ pairs within 2Å after superposition were included in the matrix calculation.

### Carbamate trapping

Purified hCx26 (0.7 mg) in stabilising buffer (100 mM NaCl, 1.25 mM NaH_2_PO_4_, 3 mM KCl, 10 mM D-glucose, 1 mM MgSO_4_, 2 mM CaCl_2_, 0.03 % (w/v) *n*-Dodecyl β-D-maltoside, 1 mM DTT, pH 7.5) was incubated with NaHCO_3_ (50 mM, equivalent to 90 mmHg PCO_2_ for hypercapnic conditions or 15 mM, equivalent to 20 mmHg PCO_2_ for hypocapnic conditions) and added to a pH stat (Titrando 902, Metrohm) equipped with pH probe and burette. TEO (280 mg, Sigma UK) was added in three step-wise increments in phosphate buffer (50 mM, 1 mL, pH 7.4). The reaction pH was maintained at 7.4 with the addition of 1 M NaOH throughout, the pH was stabilised until reaction completion (60 min). The trapped protein was dialysed overnight (1 L, dH_2_O, 4 °C).

### Tryptic digestion

After dialysis the trapped sample supernatant was removed using vacuum centrifugation. The protein was digested using the S-trap™ method (Profiti) according to the manufacturer’s instructions as follows. The sample was resuspended in SDS lysis buffer (50 µL) and reduced with DTT (final concentration 20 mM, 95 °C, 10 min). The sample was then alkylated with iodoacetamide (final concentration 40 mM, in the dark, 30 min, room temperature). The sample was then centrifuged for 8 min at 13000, 12% (v/v) phosphoric acid (final concentration 1.2%) was added and the solution diluted with S-trap binding buffer (90% aqueous methanol, 100 mM triethylammonium bicarbonate, pH 7.1). Protein sample (300 µg) was added to an S-trap mini column and digested with trypsin (1:25 ratio, 47 °C, 1 h). The peptides were then eluted and analysed by ESI-MS-MS.

### Mass spectrometry

Peptides were analysed by ESI-MS-MS on a Qstar Pulsar QTOF mass spectrometer (Sciex). Peptides were separated using an LC gradient from 3 to 80% (v/v) acetonitrile and injected online to the mass spectrometer (IDA mode, mass range 300–1600 Da, MS accumulation time 1 s, ion source voltage 2300 V, 3 MS-MS spectra per cycle, MS-MS mass range 100–1600 Da, MS-MS accumulation time 3 s). The post-ESI-MS-MS raw data files were converted into .mgf files using the freeware MSConvert^53^ provided by Proteowizard and analysed using PEAKS Studio 10.5 software^54^ including the variable modifications ethylation (28.0313 Da at D or E), carboxyethylation (72.0211 at K) oxidation (M), acetylation (*N*-terminal) and the fixed modification carbimidomethyl (C). These data were then refined using a false discovery rate (FDR) of 1 % and a PTM AScore of 20.

## Acknowledgements

We thank the Leverhulme Trust (RPG-2015-090) and MRC (MR/P010393/1) for support. We acknowledge the Midlands Regional Cryo-EM Facility, hosted at the Warwick Advanced Bioimaging Research Technology Platform, for use of the JEOL 2100Plus, and the Midlands Regional Cryo-EM Facility, hosted at Leicester Institute of Structural and Chemical Biology for use of the FEI Titan Krios G3, both supported by MRC award reference MC_PC_17136. We are grateful to Dr Kyle Morris, Dr Corinne Smith and Dr Saskia Bakker for discussion and advice and the technical support in the School of Life Sciences, University of Warwick.

## Author Contributions

The project was initiated and supervised by ND and AC. Cloning, expression, purification and grid preparation were carried out by DB. Data collection was performed by CS, TJR, DB. Data processing was done by DB and AC with guidance from TJR and CS. DB and AC refined the structures. Mass Spectrometry processing and analysis was carried out by VL and MC. AC, DB, ND wrote the manuscript with contributions from all authors.

## Data Availability

Cryo-EM density maps have been deposited in the Electron Microscopy Data Bank (EMDB) under accession numbers EMD-xxxxx (Cx26-55mmHg PCO_2_), EMD-xxxxx (Cx26-90mmHg PCO_2_), EMD-xxx (Cx26-20mmHg PCO_2_). The four classes associated with the 55mmHg CO_2_ have been deposited with accession numbers EMD-xxxxx, EMD-xxxxx, EMD-xxxxx, EMD-xxxxx. Structure models have been deposited in the RCSB Protein Data Bank under accession numbers xxx, xxx, xxx. Data from MS-MS have been deposited in the PRIDE database with accession number xxx.

## Extended Data

**Extended Data Fig. 1:**
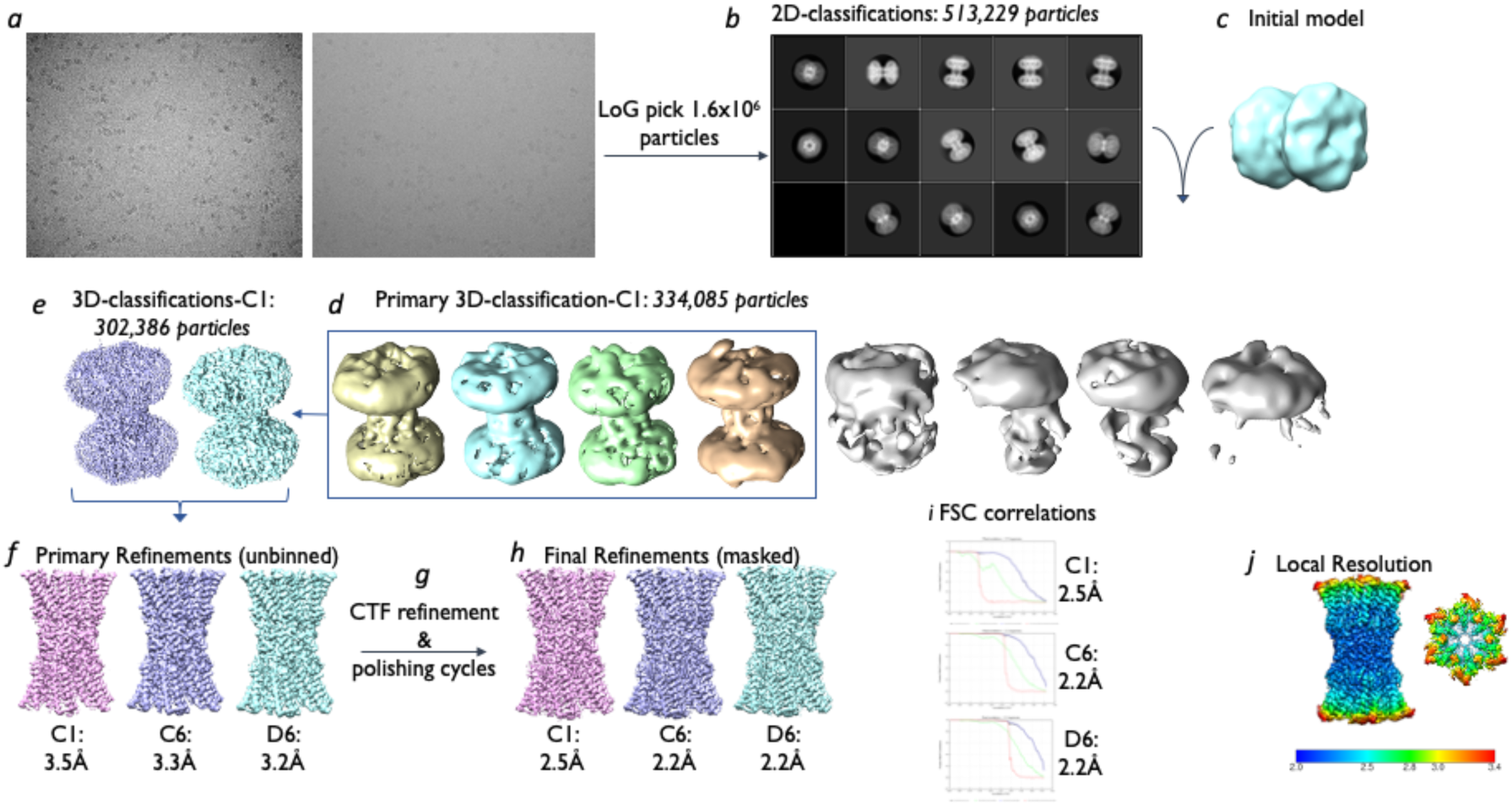
Workflow for cryo-EM data processing in Relion^38^. The resolution of reconstructions using C1, C6 and D6 symmetry is shown before and after multiple rounds of particle polishing and CTF refinement. The local resolution map is coloured according to resolution estimated in ResMap^47^.

**Extended Data Fig. 2:**
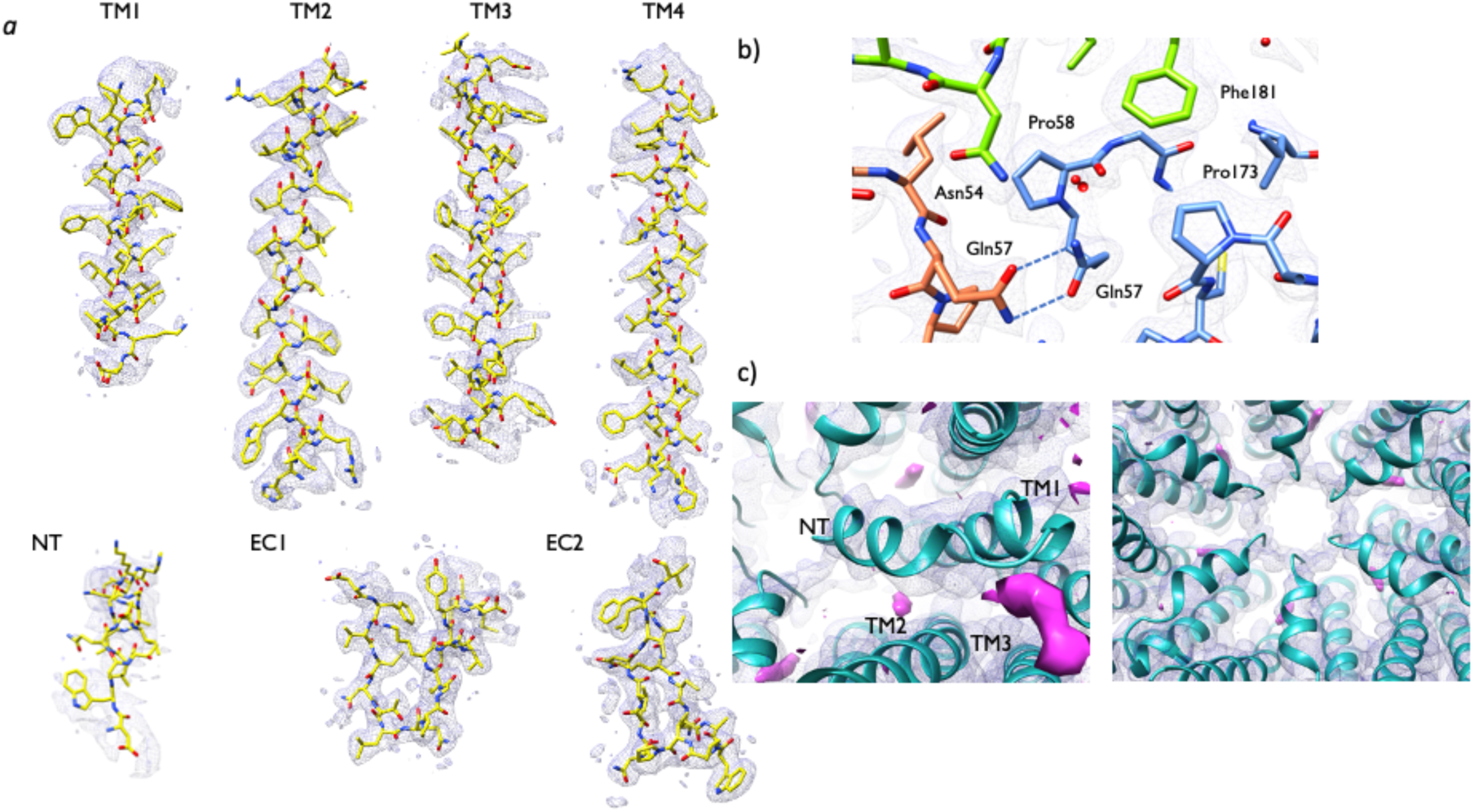
Density for 55mmHg PCO_2_ cryo-EM map. **a)** Density associated with main structural features (Transmembrane (TM) helices 1-4, the N-terminal helix (NT) and Extracellular (ET) loops 1 and 2.) No sharpening has been applied. (The N-terminus has been shown with a lower threshold (0.0117 *vs* 0.0126).) **b)** Electron density associated with the gap junction with a map sharpened in Relion (Sharpening B −22Å^2^). **c)** Features in the density maps that could not be interpreted (shown in pink). A sausage shaped density was observed in the middle of the pore. There was also density extending from TM3 to the N-terminal helix. The density associated with the N-terminus can be seen to form a ring in the centre of the pore.

**Extended Data Fig. 3:**
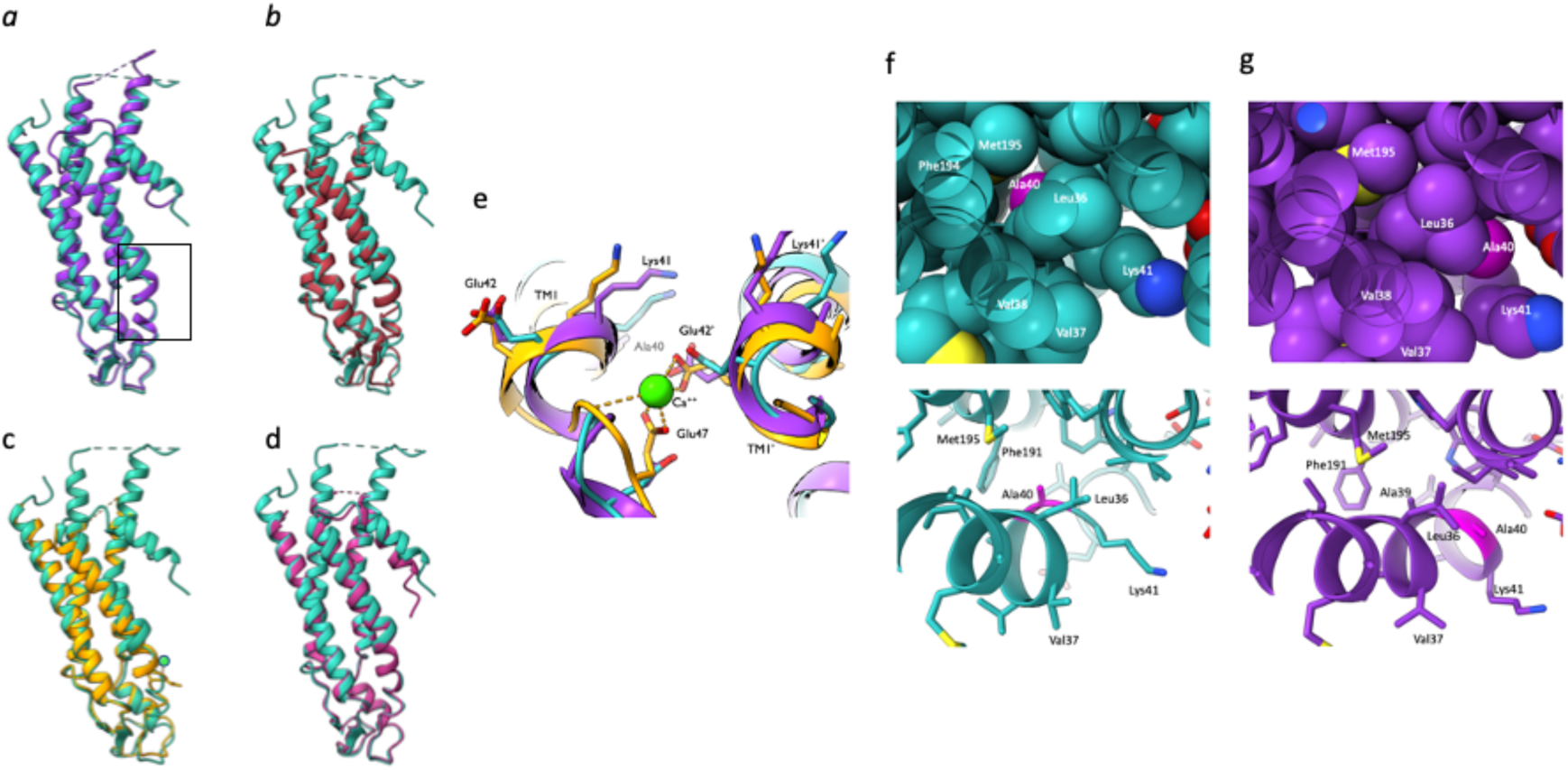
Comparison of N-terminus and TM1 in structures of connexins. Structure of subunit of Cx26-EM structure (cyan) superposed on crystal structures of Cx26 **a)** 2ZW3^33^ (purple): crystallised in UDM **b)** 5ER7^34^ (orange): calcium-bound, crystallised in a facial amphiphile **c)** 5ERA^34^ (brown) as (b) but without calcium and **d)** 6MHQ^35^ (violet) the cryo-EM structure of Cx46. The N-termini are not observed in either 5ER7 or 5ERA. The region between EC1 and TM1 (boxed) varies between the Cx26 structures. **e)** Comparison around the region of Glu42. Residues between Val37 and Glu42 vary amongst all the structures; residues Glu42 to Glu47 between the calcium-bound and non-bound structures. Using the refined cryo-EM structure as a search model for molecular replacement against structure factor amplitudes associated with either the 5ER7 or 5ERA models clearly shows the resulting density to be consistent with the crystallographic models. **f-g** Comparison of position of Ala40 in cryo-EM structure relative to crystal structure **f)** Cryo-EM structure. The top view shows a sphere representation of the protein. Carbon atoms are shown in cyan, except for Ala40 in magenta. The bottom panel shows a ribbon representation for orientation. **g)** the crystal structure 2ZW3 in the same view as (f).

**Extended Data Fig. 4:**
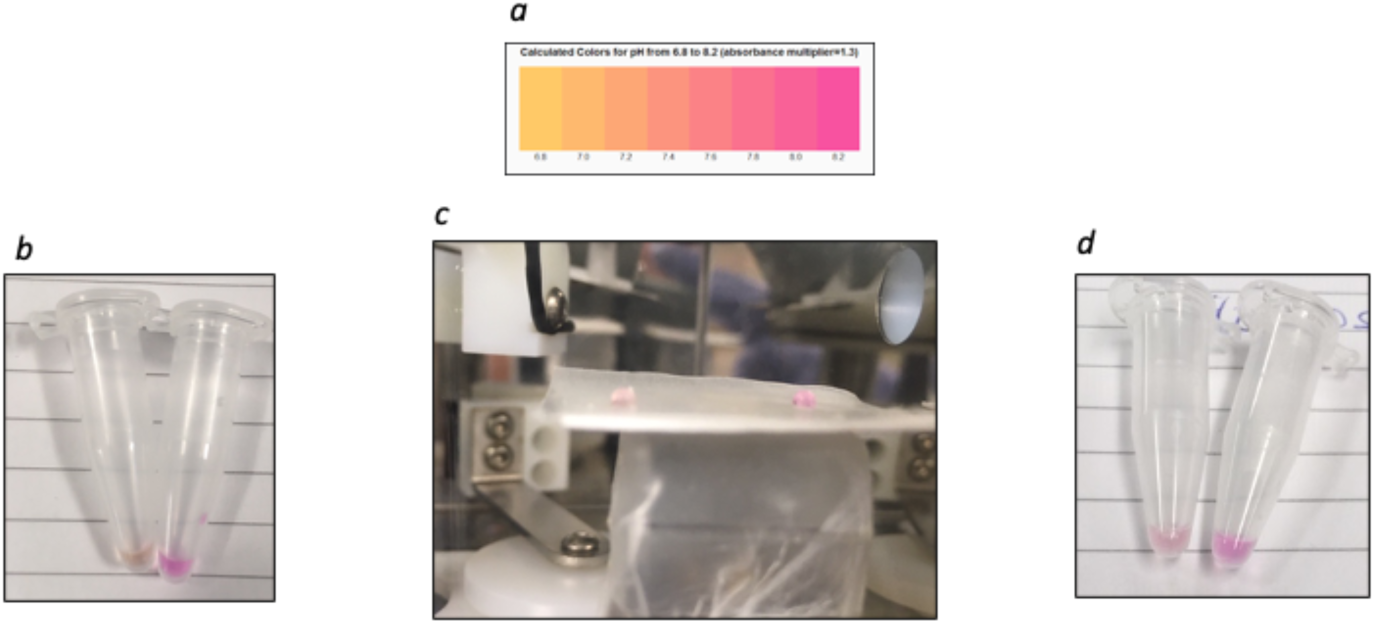
Control to ensure that the pH was the same in all samples before vitrification. **a)** Colour chart of phenol red pH indicator (dynamic range pH6.8 to pH8.2) **b)** Phenol Red (1:200 dilution) buffer gassed with CO_2_ (left) against buffer without gassing (right) **c)** 3μl drop of buffer gassed (left) and ungassed (right) in the plunge freezer chamber after 30 seconds in the equilibrated atmosphere **d)** Tubes from (a) 24 hours later demonstrating the buffer holds the pH in a sealed environment.

**Extended Data Fig. 5:**
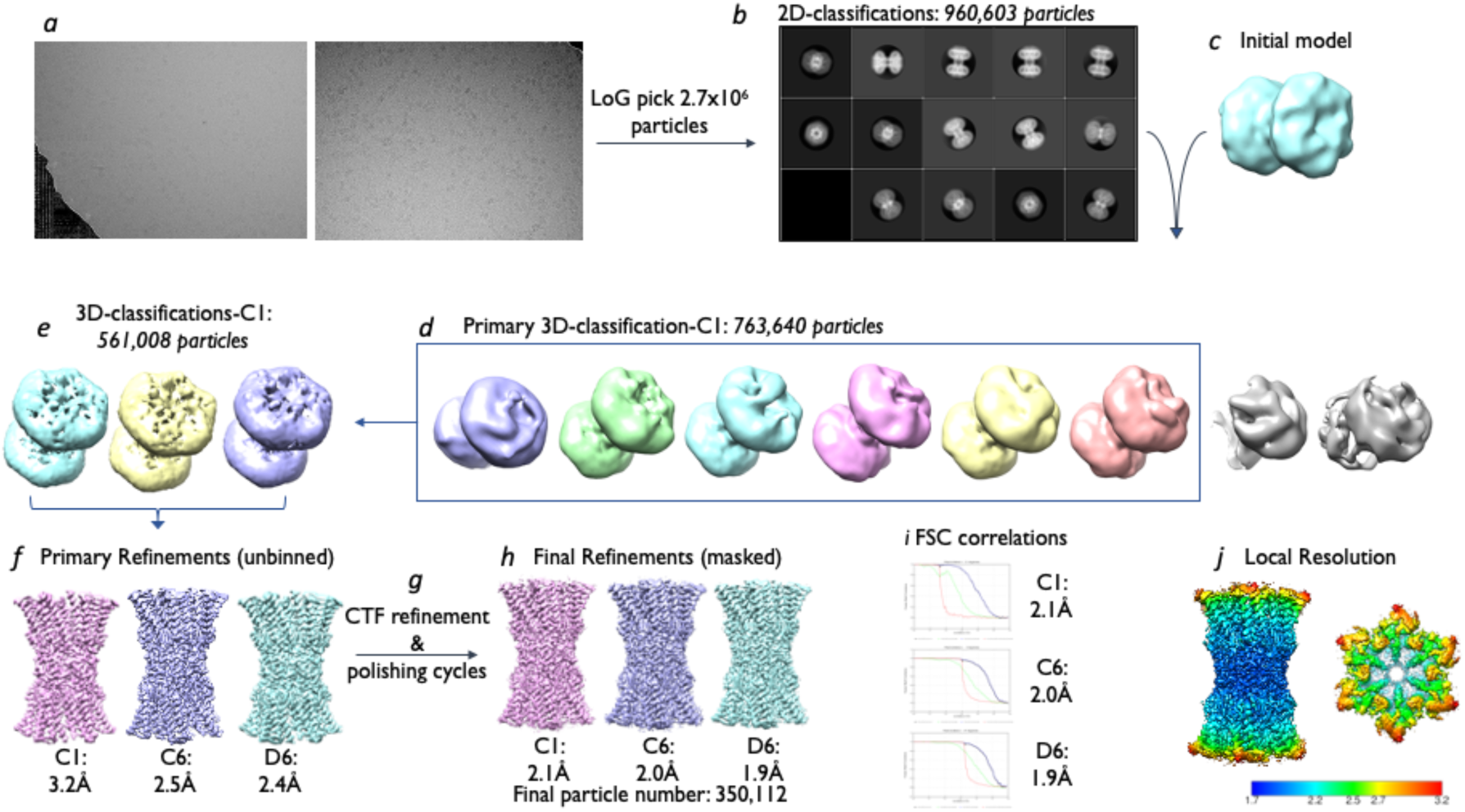
Workflow for processing of cryo-EM data associated with 90mmHg PCO_2_. As for Extended Data Fig. 1 the resolution of reconstructions using C1, C6 and D6 symmetry is shown before and after multiple rounds of particle polishing and CTF refinement. The local resolution map is coloured according to resolution estimated in ResMap^47^.

**Extended Data Fig. 6:**
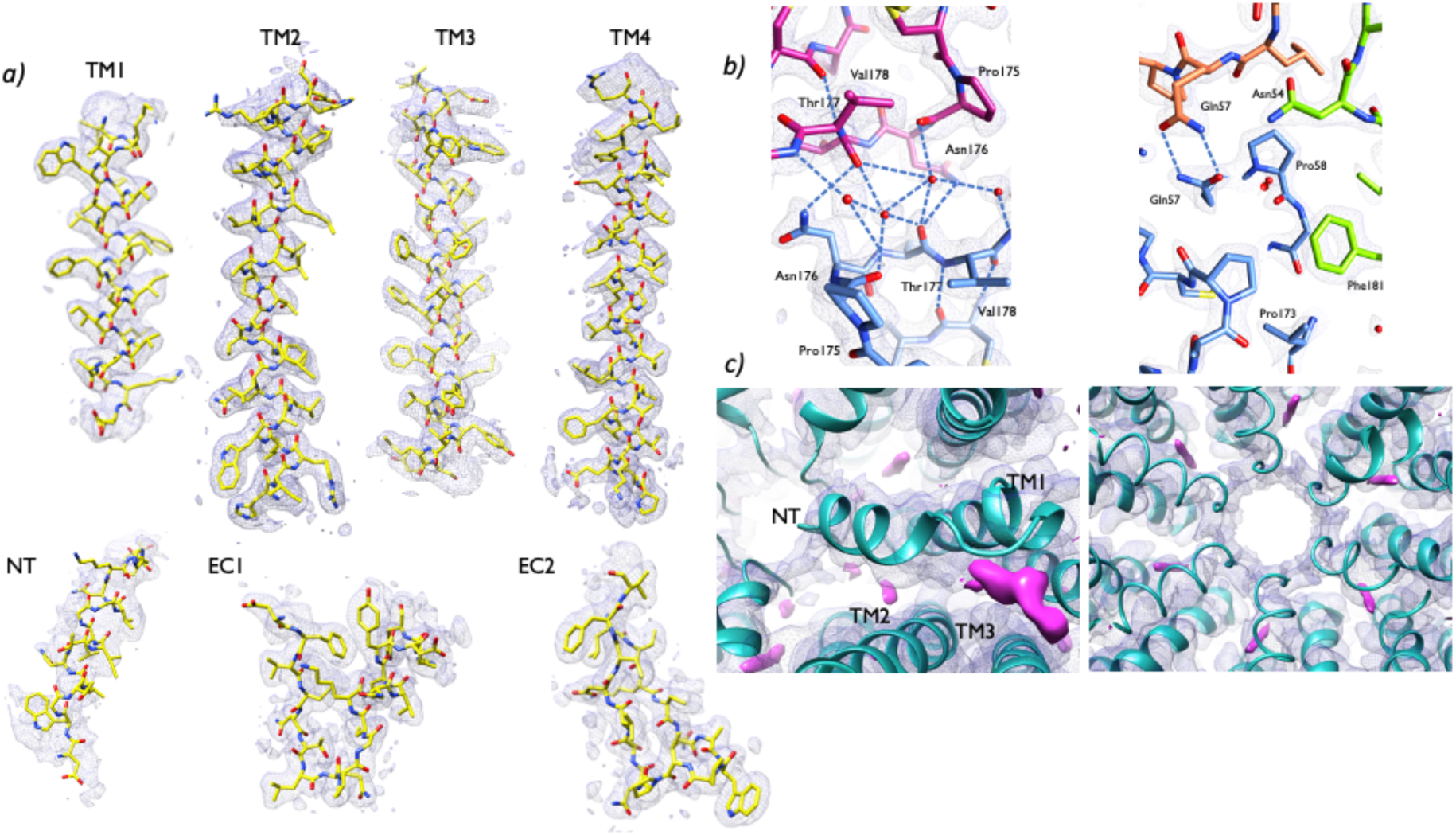
Density for 90mmHg PCO_2_ maps. Equivalent to Extended Data Fig. 2 **a)** Density associated with main structural features (Transmembrane (TM) helices 1-4, the N-terminal helix (NT) and Extracellular (ET) loops 1 and 2.) No sharpening has been applied. (The N-terminus has been shown with a lower threshold (0.0065 *vs* 0.0069).) **b)** Electron density associated with the gap junction with a map sharpened in phenix.autosharp^51^. **c)** Features in the density maps that could not be interpreted (shown in pink). A sausage shaped density was observed in the middle of the pore. There was also density extending from TM3 to the N-terminal helix. The density associated with the N-terminus can be seen to form a ring in the centre of the pore.

**Extended Data Fig. 7:**
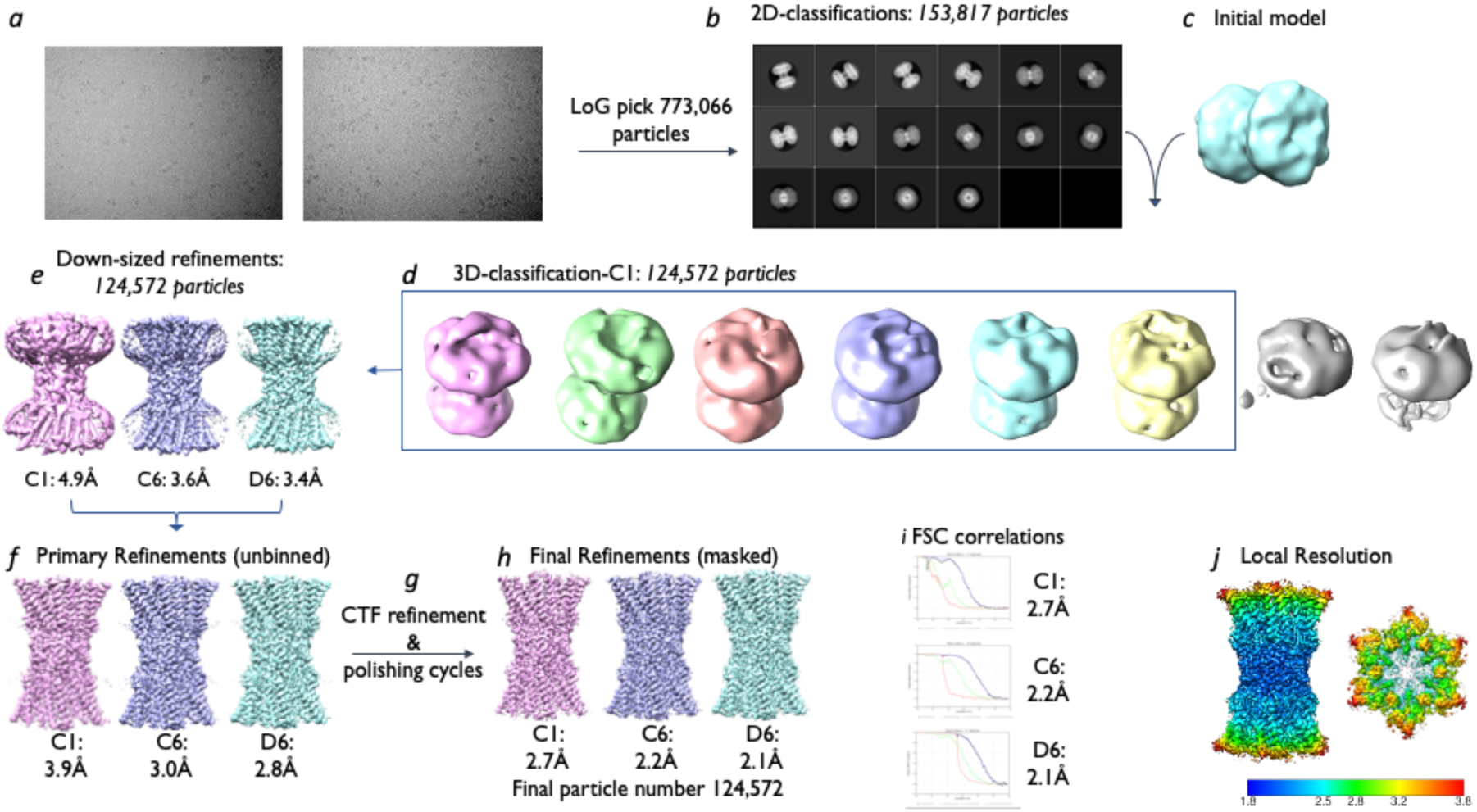
Workflow for processing of cryo-EM data associated with 20mmHg PCO_2_. As for Extended Data Fig. 1 the resolution of reconstructions using C1, C6 and D6 symmetry is shown before and after multiple rounds of particle polishing and CTF refinement. The local resolution map is coloured according to resolution estimated in ResMap^47^.

**Extended Data Fig. 8:**
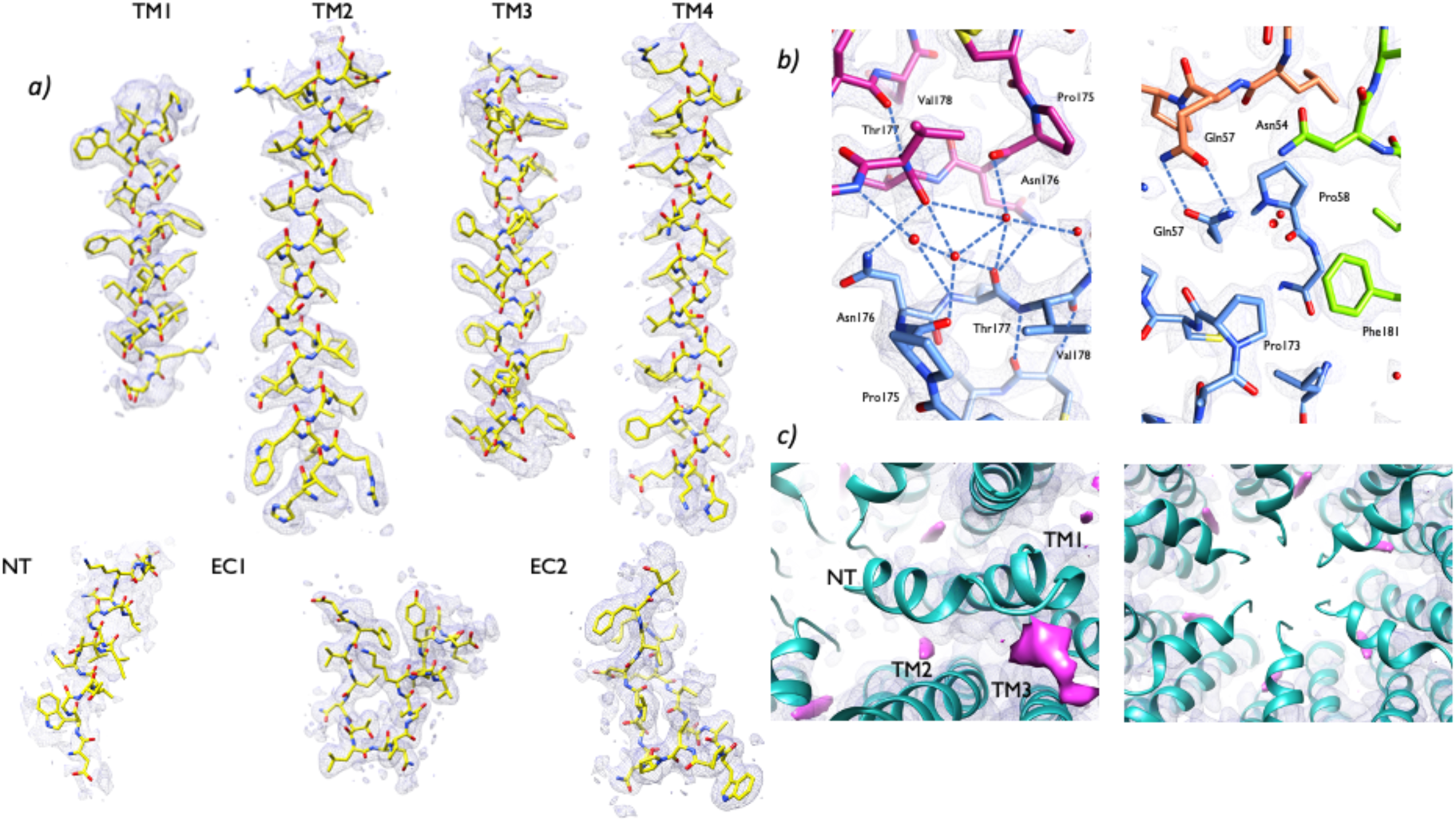
Density for 20mmHg PCO_2_ maps. Equivalent to Extended Data Fig. 2 **a)** Density associated with main structural features (Transmembrane (TM) helices 1-4, the N-terminal helix (NT) and Extracellular (ET) loops 1 and 2.) No sharpening has been applied. (The N-terminus has been shown with a lower threshold (0.0065 *vs* 0.0069).) **b)** Electron density associated with the gap junction with a map sharpened in phenix.autosharp^51^. **c)** Features in the density maps that could not be interpreted (shown in pink). A sausage shaped density was observed in the middle of the pore. There was also density extending from TM3 to the N-terminal helix. The density associated with the N-terminus is much less defined.

**Extended Data Fig. 9:**
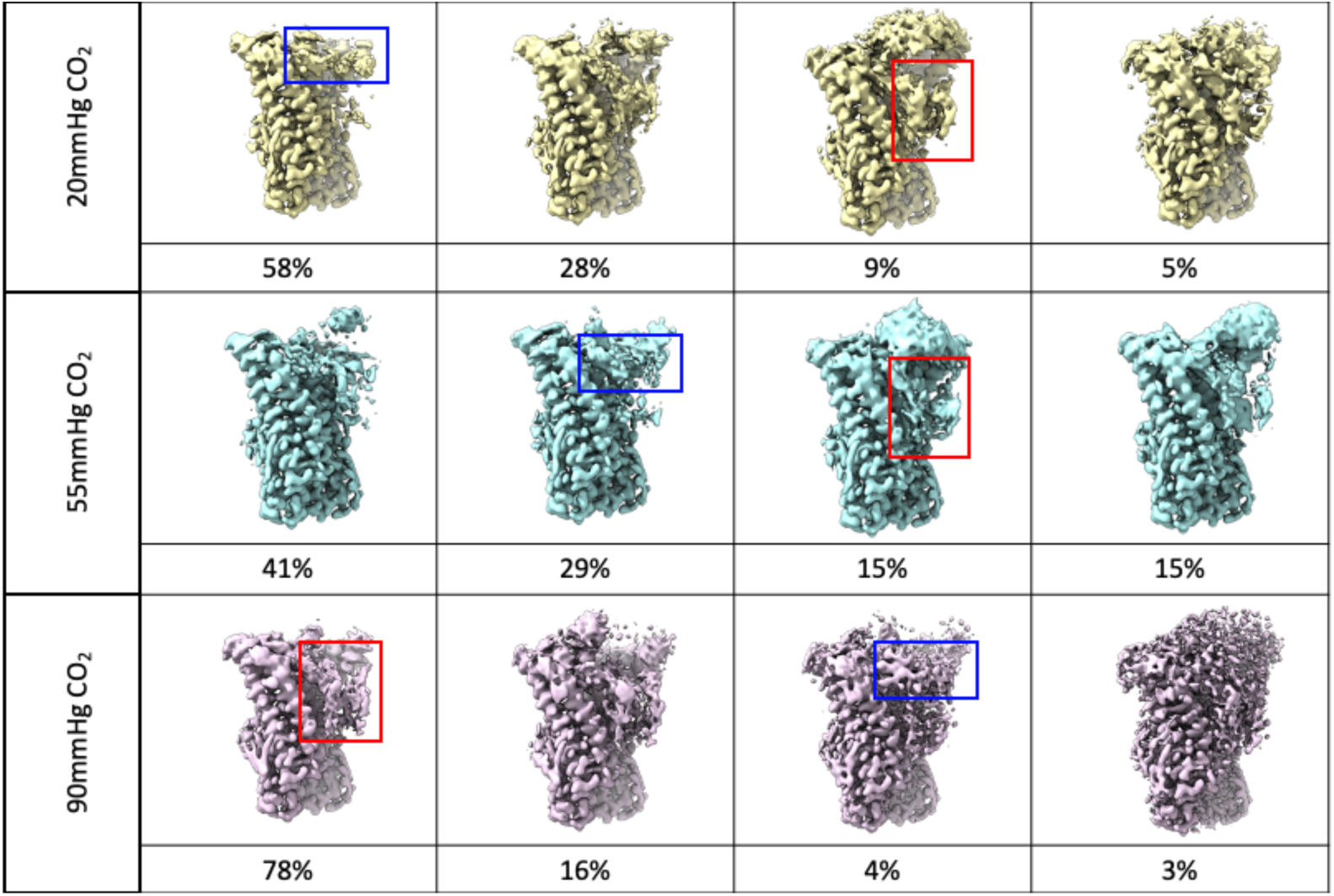
Classification as shown in Fig. 4 for the data associated with all three levels of CO_2_. From each data set 125,000 particles were randomly selected from the final D6 refinement. These were symmetry expanded in D6 and the same mask was used for particle subtraction and 3D classification, with appropriate change in pixel and box size for the 55mmHg data. The percentage of particles in each class is shown below each class. Red boxes indicate the full, extended NT position, blue boxes indicate the truncated, raised NT location. The remaining two classes in each data set represent intermediates between these two extreme classes.

**Extended Data Movie 1: Variability analysis in cryoSPARC**^37^. The variability analysis was performed at 4.5Å using a mask covering a hemichannel with particles downsampled to 1.5Å/pixel, refined in D6 and symmetry expanded. **a)** The movie comprises 20 frames from component 1 of the analysis. **b)** Each of the frames in the movie were 2-fold averaged (C2) to give a movie that shows the same general trend as in (a), but is less noisy. **c)** Side view of (b) showing the constriction formed by the N-termini. All analyses carried out, varying masks, particles subsets *etc.* showed a similar pattern of movement, which is consistent with the variability observed amongst the various connexin structures. It is noticeable that, even though symmetry expansion was applied before the analysis so that all the subunits are effectively, equivalent, the movements are not the same across the subunits. This might suggest that the subunits are not-coordinated.

**Extended Data Movie 2:** Morph between structures associated with the classification shown in Fig. 4. The morph is shown between two of the structures. As the morph only includes matching pairs the complete N-termini are not shown because this was curtailed in one of the structures. Each of the two subunits has been coloured according to the (reversed) colours of the rainbow with blue at the N-terminus to red at the C-terminus. TM2 is shown in dark green and TM3 in light green. The positions of the residues that when mutated to a bulkier residue lead to KID syndrome are shown in magenta and purple. Those in magenta affect the more flexible region of the N-terminus and TM2. Those in purple affect TM1 and the Ca^2+^ site.

**Extended Data Table 1:** Cryo-EM data collection and processing statistics.

|  | **55 mmHg CO<sub>2</sub>** | **90 mmHg CO<sub>2</sub>** | **20 mmHg CO<sub>2</sub>** |
| --- | --- | --- | --- |
| Voltage (kV) | 300 | 300 | 300 |
| Magnification (x1000) X | 81 | 105 | 105 |
| Camera | K3 | K3 | K3 |
| Camera mode | Super-resolution | Super-resolution | Super-resolution |
| Energy filter (eV) | 20 | 20 | 20 |
| Defocus range (mm) | -2 to -1 | -2 to -1 | -2 to -1 |
| Pixel size (Å/pix) | 1.09 | 0.85 | 0.85 |
| Dose on detector (e<sup>-</sup>/pix/sec) | 10 | 10 | 10 |
| Dose on sample (e<sup>-</sup>/pix/sec) | 10.9 | 11 | 11 |
| Exposure time | 5 | 3 | 3 |
| No. of images | 4003 | 11647 | 3305 |
| Frames per image | 45 | 45 | 45 |
| Initial particle number | 1679668 | 2655822 | 773121 |
| Final particle number | 344085 | 360112 | 124572 |
| **Resolution<sup>a</sup>** |  |  |  |
| masked D6 | 2.2 | 1.9 | 2.1 |
| masked C6 | 2.2 | 2.0 | 2.2 |
| masked C1 | 2.5 | 2.1 | 2.7 |

<sup>a)</sup> From Relion\_postprocess<sup>38</sup>

**Extended Data Table 2.** Cryo-EM refinement and validation statistics.

| **Refinement** | **55mm Hg CO<sub>2</sub>** | **90mm Hg CO<sub>2</sub>** | **20mmHg CO<sub>2</sub>** |
| --- | --- | --- | --- |
| Initial model used (PDB code) | 2ZW3 | 55mm Hg CO<sub>2</sub> | 55mm Hg CO<sub>2</sub> |
| Resolution (Å; FSC=0.5) Map | 2.2 | 1.9 | 2.1 |
| Sharpening B factor (Å<sup>2</sup>) | 0 | 0 | 0 |
| **Model composition** | 12 subunits | 12 subunits | 12 subunits |
| Non-hydrogen atoms | 19097 | 19313 | 19205 |
| Protein residues | 2244 | 2268 | 2256 |
| Water | 305 | 305 | 305 |
| Ligand: lipid/detergent | 60 | 60 | 60 |
| **B factor (Å<sup>2</sup>)** |  |  |  |
| protein | 71 | 63 | 72 |
| water | 42 | 36 | 42 |
| Lipid/detergent | 65 | 57 | 64 |
| **R.m.s. deviations** |  |  |  |
| Bond lengths (Å) | 0.008 | 0.002 | 0.004 |
| Bond angles (°) | 0.615 | 0.479 | 0.501 |
| **Validation** |  |  |  |
| MolProbity score | 1.45 | 1.05 | 1.29 |
| Clashscore | 3.04 | 2.19 | 1.99 |
| Rotamer outliers (%) | 2.94 | 1.18 | 2.38 |
| **Ramachandran plot** |  |  |  |
| Favoured (%) | 98.36 | 98.92 | 97.83 |
| Allowed (%) | 1.64 | 1.08 | 2.17 |
| Disallowed (%) | 0.0 | 0.0 | 0.0 |
| **Map resolution (Å)** | 2.2 | 1.9 | 2.1 |
| FSC threshold | 0.143 | 0.143 | 0.143 |
| Map resolution range (Å) (unmasked) | 2.1/2.2/2.2 | 1.8/1.8/2.0 | 1.9/1.9/2.2 |
| Map resolution range (Å) (masked) | 2.1/2.1/2.2 | 1.7/1.8/1.9 | 1.9/1.9/2.1 |
| **Correlation coefficients** |  |  |  |
| CC (mask) | 0.88 | 0.90 | 0.90 |
| CC (box) | 0.84 | 0.82 | 0.82 |
| CC (peaks) | 0.76 | 0.78 | 0.76 |
| CC (volume) | 0.87 | 0.89 | 0.89 |
| Mean CC for ligands | 0.80 | 0.85 | 0.80 |

**Extended Data Table 3:** Refinement and Validation Statistics 55mmHg PCO_2_ dimer models.

|  | **Class 1** | **Class 2** | **Class 3** | **Class 4** |
| --- | --- | --- | --- | --- |
| Initial model used | 55mmHg<br>CO<sub>2</sub> | 55mmHg<br>CO<sub>2</sub> | 55mmHg<br>CO<sub>2</sub> | 55mmHg<br>CO<sub>2</sub> |
| Resolution (Å) | 2.6 | 2.7 | 2.9 | 2.9 |
| Sharpening B factor (Å<sup>2</sup>) | 0 | 0 | 0 | 0 |
| Model composition | 2 subunits | 2 subunits | 2 subunits | 2 subunits |
| Non-hydrogen atoms | 3076 | 3002 | 3056 | 3204 |
| Protein residues | 366 | 359 | 368 | 383 |
| Water | 14 | 7 | 4 | 8 |
| Ligand: lipid/detergent | 10 | 6 | 6 | 8 |
| **B factor (Å<sup>2</sup>)** |  |  |  |  |
| protein | 108 | 102 | 92 | 107 |
| water | 83 | 83 | 69 | 87 |
| Lipid/detergent | 99 | 88 | 77 | 90 |
| **R.m.s. deviations** |  |  |  |  |
| Bond lengths (Å) | 0.004 | 0.007 | 0.006 | 0.008 |
| Bond angles (°) | 0.586 | 0.627 | 0.629 | 0.692 |
| **Validation** |  |  |  |  |
| MolProbity score | 1.40 | 1.25 | 1.56 | 1.62 |
| Clashscore | 3.73 | 3.49 | 5.39 | 5.92 |
| Rotamer outliers (%) | 0 | 0 | 0 | 0 |
| **Ramachandran plot** |  |  |  |  |
| Favoured (%) | 96.37 | 97.44 | 96.11 | 95.73 |
| Allowed (%) | 3.63 | 2.56 | 3.89 | 4.27 |
| Disallowed (%) | 0.0 | 0.0 | 0.0 | 0.0 |
| **Map resolution (Å)** | 2.6 | 2.7 | 2.9 | 2.9 |
| FSC threshold | 0.143 | 0.143 | 0.143 | 0.143 |
| Map resolution range (Å)<br>(unmasked) | 2.4/2.6/2.9 | 2.4/2.7/2.9 | 2.6/2.8/3.0 | 2.6/2.8/3.0 |
| Map resolution range (Å)<br>(masked) | 2.4/2.6/2.8 | 2.4/2.6/2.9 | 2.6/2.7/3.0 | 2.4/2.7/3.0 |
| **Correlation coefficients** | 0.85 | 0.87 | 0.88 | 0.86 |
| CC (mask) | 0.77 | 0.78 | 0.78 | 0.79 |
| CC (box) | 0.63 | 0.66 | 0.66 | 0.67 |
| CC (peaks) | 0.84 | 0.86 | 0.86 | 0.85 |
| CC (volume) | 0.68 | 0.70 | 0.70 | 0.69 |

## Notes

### Competing Interest Statement

The authors have declared no competing interest.

